# Neighborhood-Informed Positional Information for Precise Cell Identity Specification

**DOI:** 10.1101/2025.03.17.643609

**Authors:** Michal Erez, Roy Friedman, Mor Nitzan

**Affiliations:** School of Computer Science and Engineering, The Hebrew University, Jerusalem, Israel; Racah Institute of Physics, The Hebrew University, Jerusalem, Israel; Faculty of Medicine, The Hebrew University, Jerusalem, Israel

**Author notes:** Equal contribution (Michal Erez, Roy Friedman).

## Abstract

During development cells reliably establish their identities, a process that is enabled in part by positional information encoded in gene expression patterns. Previous works showed that cells in *Drosophila* embryos can utilize this information to decode their position along the anterior-posterior axis with 1% accuracy. However, this precision is insufficient to uniquely determine position, leading to a positional information gap. Here, we propose a neighborhood-informed information-theoretic framework where cells integrate local gene expression information as well as information from neighboring cells. We formulate how much additional information exists in neighboring cells as a function of spatial variation in gene expression. In *Drosophila* embryos, we show that the additional information encoded by local neighborhoods is sufficient to uniquely specify cell identities, closing the information gap. Furthermore, neighborhood-informed decoders predict cell positions and downstream gene expression patterns more accurately than cell-independent decoders, resulting in lower decoding variability, which is maintained in mutant embryos. Our results provide a basis for the analysis of cellular decision making in the context of their microenvironments.

## I. Introduction

Multicellular self-organization during development is an incredibly precise and reproducible process [1–5]. In several developmental contexts, the precision of self-organization is derived from positional information, where extrinsic signals, such as morphogens, carry information that enables cells to effectively decipher their position within an embryo or within a tissue. This positional information, in turn, can be used by cells to specify their identity as a function of their position [6–11].

In recent years, a seminal line of works formalized an information-theoretic framework for positional information, as well as its accuracy and limitations, with a focus on positional information of individual cells in the well-studied model of the *Drosophila* embryo [9, 10, 12, 13]. The developmental dynamics in *Drosophila* embryos, where gene expression patterns and the different stages of development have largely been mapped out [14–16], provide a useful test-case for the study of positional information [8–10, 12, 13, 17]. In these works, positional information was studied along the main body axis of the embryo, the anterior-posterior (AP) axis, where information from maternal morphogens provides input to an interacting network of *gap genes* {knirps (kni), Krüppel (Kr), hunchback (Hb), giant (Gt)}, whose expression in turn provides input to the patterned striped expression of *pair-rule genes* {Eve (eve), paired (prd), runt (run)}. The pair-rule genes, thereby, map the segmented body plan of the developed fly. Given this flow of information, it was suggested that expression patterns of gap genes encode positional information, and expression of pair-rule genes can be considered as the positional readout of the cells [18, 19].

Indeed, gap gene expression provides individual cells in the *Drosophila* embryo enough information to decode their position along the AP axis with high precision [8, 10]. However, this *cell-independent* decoding is insufficient to fully specify unique cell positions [8, 9]. In other words, there seems to be an *information gap* between the information available to individual cells and the information required for unique cell localization [9]. It was suggested that the information gap can be theoretically bridged by long-range gene expression correlations along the AP axis [9]. This is because such gene expression correlations introduce effective constraints: they reduce the number of possible collective (AP axis wide) gene expression states, and thus, reduce the amount of information needed for each cell to specify its position. It remains unclear, however, how cells can directly utilize this information to specify their position.

Here, we suggest a formulation for positional information encoding that includes cell-cell interactions (whether direct or indirect) via a *neighborhood-informed* framework, where each cell can aggregate information provided by both external signals as well as information from its cellular microenvironment. We then show that such *neighborhood-informed* positional decoding can directly close the information gap, namely, that it allows cells to uniquely decode their position along the AP axis.

First, we show that the long-range correlations mentioned above can be viewed as a consequence of correlations in gene expression between neighboring cells. These local expression correlations completely dictate long-range correlations discussed in previous work [9], and above a certain threshold can close the information gap. Empirically, we find that these pairwise correlations are indeed higher than the theoretical bound, based on measurements of gene expression along the AP axis in *Drosophila* embryos [8] (Supplementary Information). This result indicates that the information missing for cells to uniquely specify their position exists in their local neighborhood, but does not explain how this collectively encoded positional information can be directly utilized by individual cells to infer their position. We address this via a *neighborhood-informed* positional information framework. In our framework, the amount of positional information encoded in local expression levels can be inferred from the performance of the optimal probabilistic position decoder given the gene expression in a certain position and its microenvironment. Formally, the neighborhood-informed probabilistic position decoder is defined as 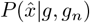 where 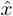 is the predicted position given the gene expression at a specific position, *g*, and that of its neighborhood, *g*_*n*_. This is in contrast to a cell-independent view, where the decoder only has access to *g*. The information that can be extracted in such a case is inversely proportional to the variance in the prediction of the position [10]. Through this framework, we are able to mathematically formulate how much additional positional information can be gained by incorporating information from neighbors. We find that the variance of the positional prediction using a neighborhood-informed decoder is consistently low enough to close the gap in positional information, unlike the cell-independent decoder [8]. Furthermore, we demonstrate that the neighborhood-informed decoded positions can better predict pair-rule gene expression profiles.

Finally, we test our framework in the context of mutant *Drosophila* embryos, where the maternal-input has been perturbed [8]. Pair-rule gene expression in such mutants, similarly to wild-type embryos, is approximately determined by gap gene expression profiles [8]. We re-evaluate the position decoding in these maternal-input perturbed embryos to include neighborhood information, assuming an encoder that was optimized in wild-type embryos. In this case as well, the neighborhood-informed decoder maps gap gene expression to pair-rule gene expression with higher positional certainty than a cell-independent decoder. These results provide additional support that cells may utilize neighborhood information directly to infer cell state, such as position, and provide a framework to test neighborhood-informed positional information encoding in diverse biological contexts.

## II. Results

### A. Short-range gene expression correlations close the positional information gap in the *Drosophila* embryo

To estimate the amount of positional information encoded in gap gene expression, we analyze a position predictor given gap gene expression levels. Mathematically, the decoded position 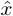 depends on observed gap gene expression levels, *g*, at position *x*. Under the assumption that 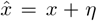 where *η* is Gaussian distributed, then the amount of information that the expression levels of gap genes contain regarding the true position is inversely related to the position estimate error, namely the variance 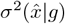 of the predicted location. Concretely, the amount of positional information *I*_position_ that can be extracted from gap gene expression levels along the AP axis can be formulated as [9]:

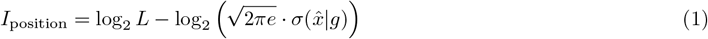

where *L* is the length of the embryo. It was previously shown [10, 13] that there exists enough information in the local expression levels of gap genes along the AP axis to estimate the position of a cell with an accuracy of approximately 1% of the embryo’s length. While 1% precision sounds extremely accurate, it is insufficient for the task of uniquely determining cell position [8, 9]. This lack of information is encapsulated in the variance of the position estimate and is formalized through the *information gap* [9]:

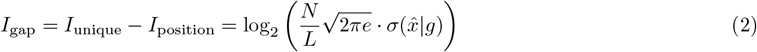

where *I*_unique_ is the amount of information required to uniquely identify cellular position and *N* is the number of nuclei along the AP axis. As long as *I*_gap_ is greater than 0, it is possible, for example, for the predicted ordering of two adjacent cells to flip.

The functional forms of the positional information and information gap above do not incorporate any assumptions on the structure of gap gene expression along the AP axis. In practice, however, these gene expression patterns vary smoothly as a function of position and there exist long-range expression correlations along the AP axis [9]. This correlation structure implies that genes in adjacent positions cannot have arbitrarily different expression levels relative to each other, effectively reducing the space of possible spatial expression patterns. This constrained space of possible states means that less information is required per cell to determine its position.

Formally, McGough *et al*. modeled these long-range correlations with an exponential decay:

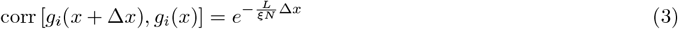

where Δ*x* is the distance between two positions along the AP axis and *g*_*i*_(*x*) is the expression level of gene *i* at position *x*. The information gap is closed when 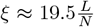 [9]. However, beyond long-range correlations, the spatial smoothness of gap gene expression implies high correlations between adjacent positions, suggesting that these longrange correlations may be rooted in pairwise correlations. This would suggest that additional positional information is encoded in local neighborhoods. We will now formalize this notion.

To simplify notation, we will assume that *L* = *N* and that the *N* positions are equally spaced along the AP axis. The correlation structure can be recapitulated when it is assumed that the gene expression levels at adjacent positions *x* and *x* + 1 are jointly Gaussian and when they have the following Markovian property (Supplementary Information):

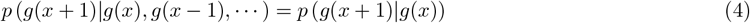

wherein the distribution of *g*(*x* + 1) is independent of all other preceding values given *g*(*x*). Under these simplifying assumptions, it can be shown that for each gene independently (Supplementary Information):

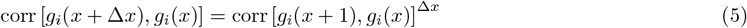

where Δ*x* is the distance between two positions along the AP axis. This means that correlations between positions at a distance of Δ*x* apart are entirely determined by the correlation between neighboring positions. Furthermore, as long as pairwise correlations are positive, there exists *ξ* such that (see Supplementary Information):

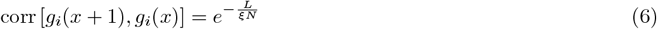

and thus we recover the long-range correlations as described by Equation 3, as arising from pairwise-correlations (Supplementary Information). As mentioned above, the positional information gap is closed when *ξ* is above some threshold, which can be translated to a threshold on the pairwise correlation:

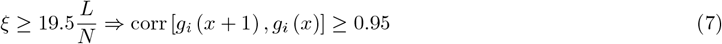

We will additionally note that the value of pairwise correlation must also be lower than 1, since otherwise there would be no additional positional information in the neighborhood due to its invariance along the AP axis.

When analyzing the wild-type (WT) *Drosophila* embryo gap gene expression profiles measured in [8], we find that throughout the AP axis, the pairwise correlations in expression of both gap and pair-rule genes are indeed within the range needed to close the information gap (Equation 7; Figure 1b,c).

**Fig. 1:**
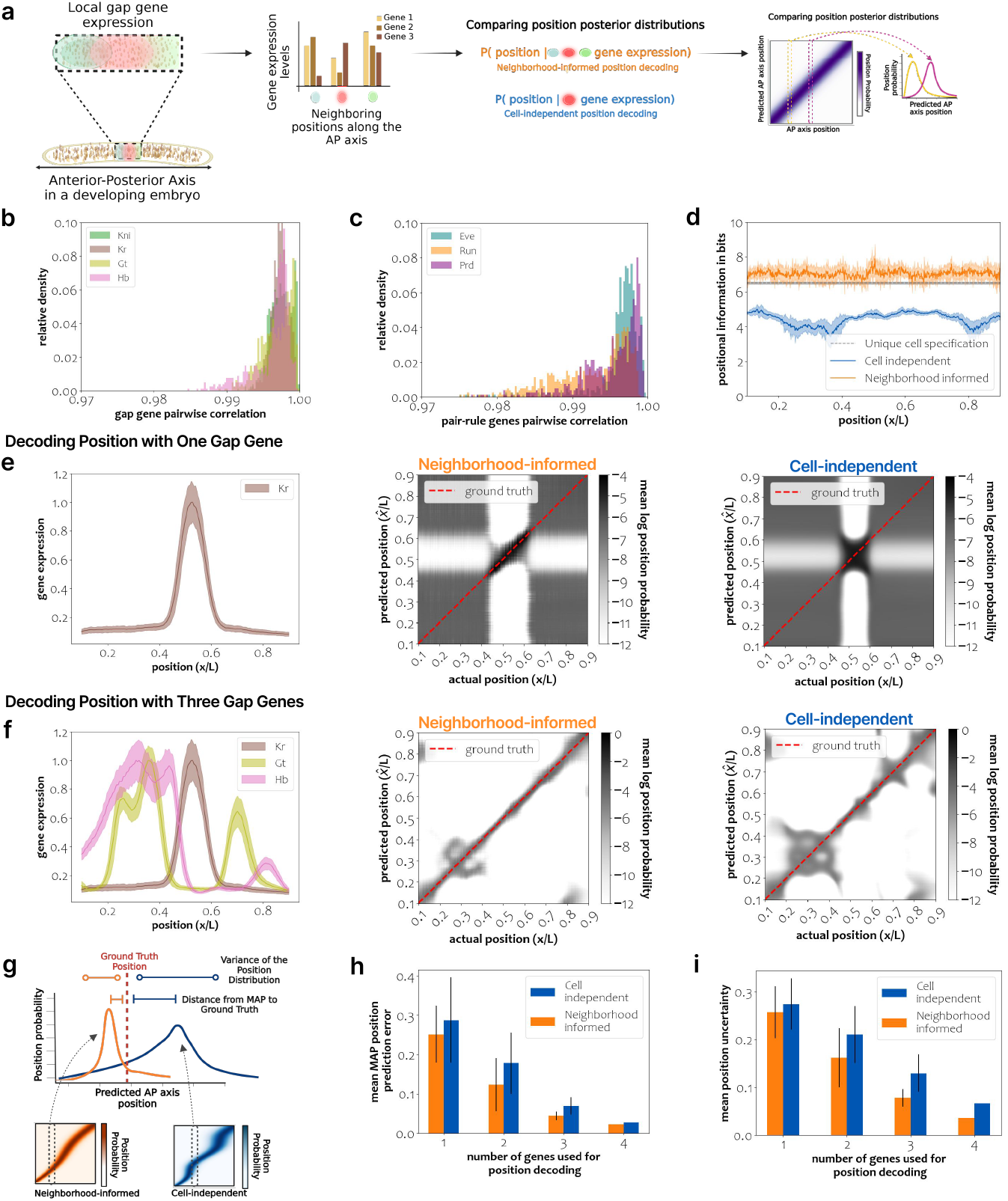
Positional Information Encoded in neighboring positions along the AP axis in *Drosophila* embryos. *(a)* A schematic diagram showing the general framework. Using gap gene expression at adjacent positions along the AP axis, we decode position in a neighborhood-informed manner and compare to cell-independent decoding. Decoding position results in positional distributions that can be represented as decoding maps (right). *(b)* The distribution of pair-wise correlation in gap gene expression of adjacent positions for each of the four gap genes, Kni, Kr, Gt, and Hb *(c)* The distribution of pair-wise correlation in expression of adjacent positions for each of the three pair-rule genes: Eve, Prd, and Run… *(d)* The positional information in bits along the AP axis given a neighborhood-informed decoding (orange) as opposed to cell-independent decoding (blue). The gray dashed line represents the information needed for unique cell identification. *(e)* (left) gap gene expression of Kr along the AP axis; neighborhood-informed (middle) and cell-independent (right) position decoding maps based on Kr gene expression. *(f)* (left) gap gene expression of Kr, Gt, Hb along the AP axis; neighborhood-informed (middle) and cell-independent (right) position decoding maps based on the expression of these three genes. *(g)* Schematic diagram portraying the comparison between the neighborhood-informed versus cell-independent position decoding distributions. *(h)* The mean error of position prediction using the MAP estimate, grouped by the number of gap genes used for decoding. For each group, we compare neighborhood-informed to cell-independent decoding. The predicted error is significantly lower for neighborhood-informed decoding (t-test p-value *<* 0.05 for embryo-to-embryo comparisons) with 2,3, and 4 genes. Decoding with one gene results in a statistically significant (t-test p-value *<* 0.05) improvement in 75% of embryos. *(i)* The standard deviation of the predicted position distribution, grouped by the number of gap genes used for decoding position, is significantly lower for neighborhood-informed decoding (t-test p-value *<* 0.05 for embryo-to-embryo comparisons for all gene subsets). Figure panels (a)and (d) were created with BioRender.com [20].

### B. Neighborhood information reduces positional error

In the previous section, we have established that at the embryo level, the combination of positional information in each cell along with the positional information encoded in short-range, pairwise gene expression correlations allows for unique cell localization. Effectively, such short-range expression correlations constrain the potential space of spatial expression patterns, which as a result, reduces the extent of positional information required by each cell to uniquely specify its position. However, it is still unclear how each cell utilizes such information and directly decodes its position. To address this, we formulate the encoding of positional information by explicitly incorporating information about neighboring gene expression, and analyze the resulting variance in the predicted position 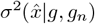 (see Methods), where *g*_*n*_ represents gene expression levels in a cell’s neighborhood.

Specifically, the neighborhood-informed position estimate error can be characterized (analogously to the cell-independent position estimate error [10]) by:

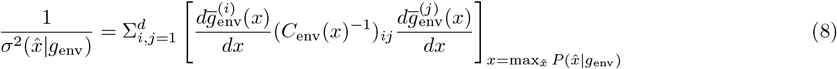

where *g*_env_ = (*g, g*_*n*_), *d* are all the gene expressions considered in 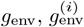 is the expression level of *i*-th gene, 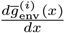 is the mean derivative of the expression level as a function of *x*, and *C*_env_(*x*) is the covariance matrix of the gene expressions in *g*_env_.

Therefore, additional information incorporated from neighboring positions is not expected to increase the error. In fact, we show that the position estimate error strictly decreases when adding neighboring information (Supplementary Information):

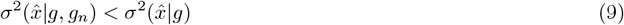

In other words, the neighborhood-informed variance of estimated position (left term in Equation 9) is bounded from above by the cell-independent variance of estimated position (right term). How to integrate neighboring information is explained in further details in Methods.

Under mild Gaussian assumptions, it can be shown that 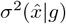 and 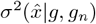 are equal to their Cramer-Rao lower bounds and Equation 9 can be written in closed form (Supplementary Information):

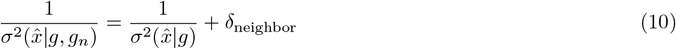

where *δ*_neighbor_ is a non-negative value related to the covariance between adjacent positions. Thus, when decoding positional information in a neighborhood-informed manner, the information gap (Equation 2) is closed when (Supplementary Information):

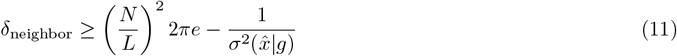

We empirically demonstrate, based on expression data in wild-type *Drosophila* embryos (Supplementary Information), that the positional information in the neighborhood-informed decoder is sufficiently high to uniquely specify cell locations and close the positional information gap, unlike the cell-independent decoder (Figure 1d). Specifically, for the *n* = 38 WT embryos dataset, we find that 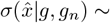 0.00195*L* when using the neighborhood-informed decoder. In comparison, the variance for the cell-independent decoder was 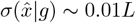, which, as stated above, is not small enough to close the information gap.

### C. Neighborhood-informed position decoding in wild-type Drosophila embryos

We will next formulate the position decoder directly, which will then enable prediction of downstream events, such as pair-rule gene expression patterns. To do so, we will first derive the full distribution 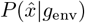 over the predicted position 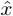 given gap gene expression *g*_env_ *≡* [*g, g*_*n*_]^*T*^ at position *x* and its neighboring environment (Figure 1a, Methods). Using Bayes’ law, the posterior distribution over predicted positions 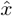 given *g*_env_ is (see Methods):

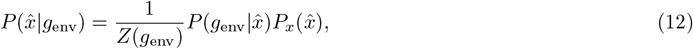

where 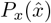 is the prior distribution over positions, 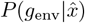 is the likelihood of *g*_env_ at position 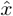, and 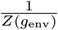 is a normalizing factor. Based on this, we can compute the posterior distribution of predicted positions, or positional distribution, given the gene expression at every position along the AP axis. When training the decoder on 254 WT embryos collected in [8] and predicting the posterior distribution for a separate dataset composed of 38 WT embryos collected in the same work, the neighborhood-informed positional distribution is more strongly peaked around the ground-truth positions relative to the cell-independent positional distribution (Figure 1e,f), showcasing the increased accuracy in maximum-a-posteriori (MAP) prediction as well as the reduction in the ambiguity over predicted positions when utilizing neighborhood information (Figure 1g-i). The results are consistent when both training and decoding of positions is performed on the same dataset (Figure **S4**).

While the positional decoding prediction error and the standard deviation of the position distribution decrease with increasing number of gap genes used for decoding for both decoder types (neighborhood-informed and cell-independent), the neighborhood-informed decoder consistently outperforms the independent decoder (Figure 1h,i).

### D. Prediction of pair-rule gene expression

Next, we will directly characterize the transition from gap gene expression to pair-rule gene expression, considered as the positional readout of the cell (Figure 2a). In other words, do gap gene expression profiles encode at the local scale the information necessary to specify pair-rule gene expression patterns?

**Fig. 2:**
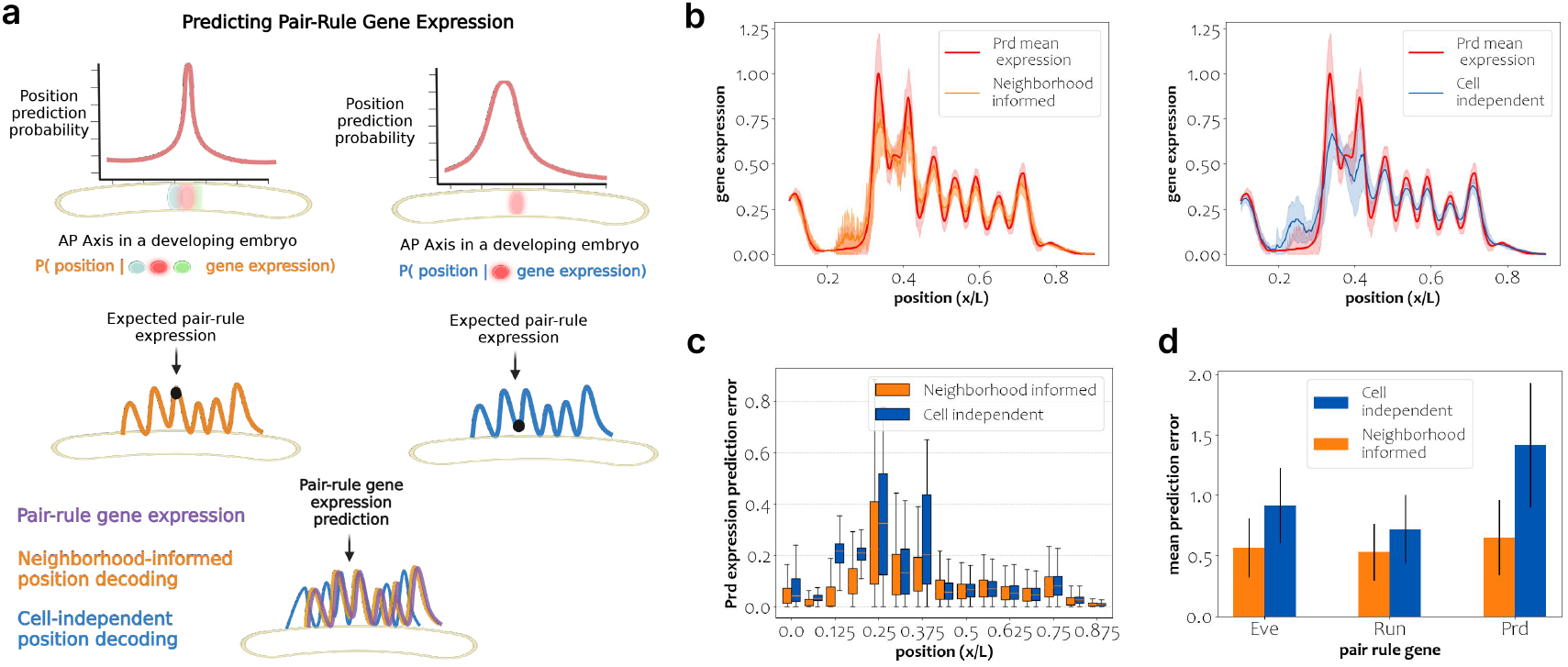
Pair-Rule stripe reconstruction in WT *Drosophila* embryos. *(a)* A schematic diagram of pair-rule gene expression prediction based on positional information. *(b)* Reconstruction of the pair rule gene Prd based on a neighborhood-informed (left) and a cell-independent (right) decoder. *(c)* Prd reconstruction error binned over 20 positions across the AP axis. *(d)* The mean reconstruction error per pair-rule gene, averaged over embryos and positions, divided by the standard deviation per position, is significantly lower for neighborhood-informed decoding (t-test p-value *<* 0.05, for all embryo-to-embryo comparisons) for Eve and Prd pair-rule expression prediction, and for 92% of embryos for Run expression prediction). Figure panel (a) was created with BioRender.com [20].

We reconstruct pair-rule gene expression profiles by utilizing the posterior distribution over predicted positions given gap gene expression. The expected pair-rule gene expression 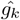 at a certain position given gap gene expression in that position and its neighborhood, *g*_env_, is then given by:

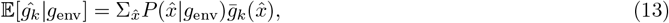

where 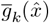 is the mean over wild-type embryos’ expression of pair-rule gene *k* at position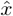. An analogous expression can be obtained for the cell-independent decoder by replacing *g*_env_ with *g*.

In Figure 2b, we show a qualitative comparison of the prediction of the expression of pair-rule gene Prd along the AP axis based on either the neighborhood-informed or cell-independent decoders, in comparison to its mean ground-truth expression (similar results for the additional pair-rule genes, Eve and Run, are shown in Figure **S1**). The neighborhood-informed prediction is more accurate, relative to the cell-independent prediction, across the AP axis (Figure 2c). This initial result generalizes, as the prediction error of pair-rule expression, averaged across all positions and wild-type embryos, is lower for the neighborhood-informed decoder, for all three pair-rule genes (Figure 2d).

### E. Decoding Position and Predicting Pair-Rule Stripes in Mutant Embryos

The empirical results from the previous sections show that substantial positional information is encoded in the local neighborhood and that an optimal decoder can decode position with higher accuracy and lower uncertainty when integrating neighborhood information. However, do cells necessarily decode position utilizing neighborhood information?

One of the strongest tools to approach such a question is by testing the given hypotheses on a perturbed system. This is possible in *Drosophila* embryos through the use of perturbed maternal signals [8], which alter the expression levels of both gap genes and pair-rule genes (Supplementary Information). Our underlying assumption, supported by previous work [8], is that although such perturbations, or mutant backgrounds, affect the spatial expression patterns of gap genes (Figure 3a), they do not substantially affect the wild-type positional decoder, and thus do not affect the mapping from gap gene expression to pair-rule gene expression, through the decoding of position (even though such decoded positions no longer reflect the actual position along the AP axis).

**Fig. 3:**
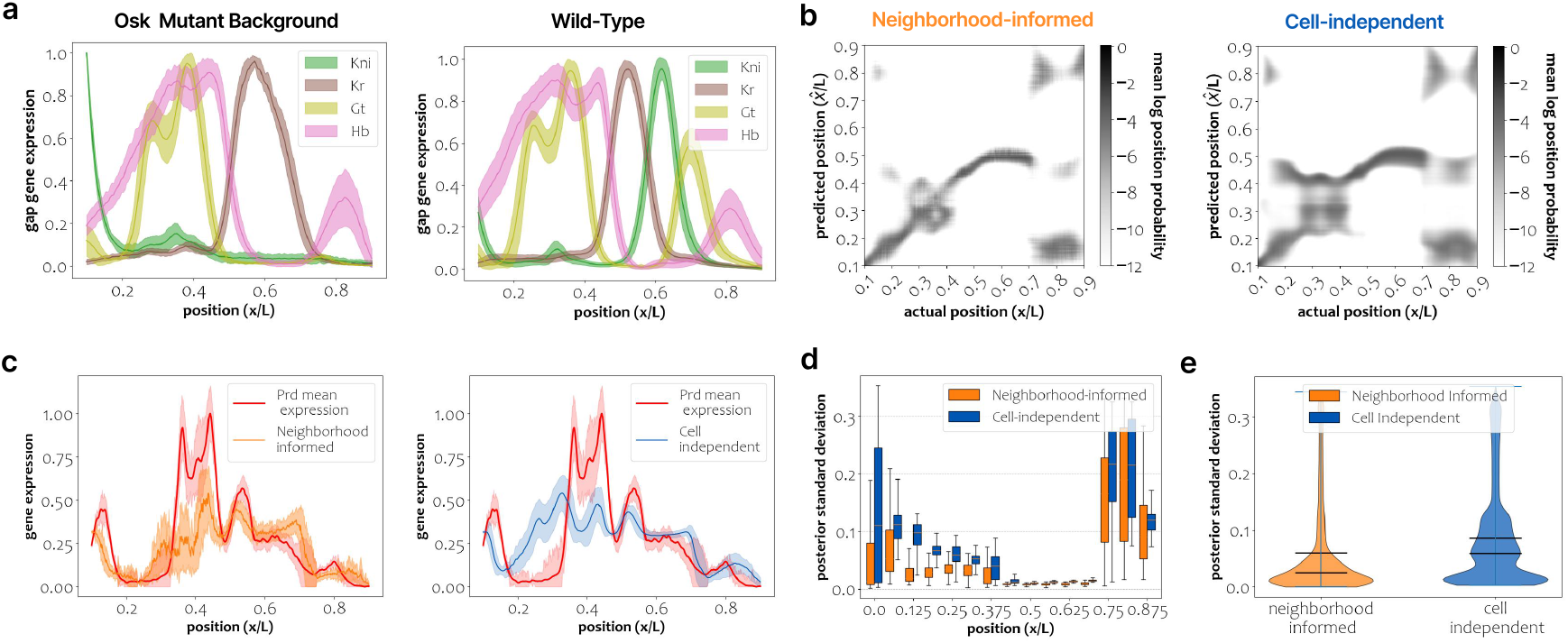
Neighborhood-informed Position Decoding and Pair-Rule Stripe Reconstruction in *osk* mutants. *(a)* Gap gene expression of all four gap genes in *osk* mutants (left) and in wild-type embryos (right). *(b)* Decoding maps given neighborhood-informed (left) and cell-independent (right) decoders. *(c)* Pair-rule expression prediction across the AP axis, using neighborhood-informed (left) and cell-independent (right) decoders. *(d)* Posterior standard deviation over predicted positions across 20 binned positions along the AP axis. *(e)* Distributions over posterior standard deviation of predicted positions (over all embryos and all positions). The Posterior standard deviation of the neighborhood-informed decoder is significantly lower than that of the cell-independent decoder (t-test p-value *<* 0.05).

Neighborhood-informed position decoding in mutant embryos results in notably less ambiguous decoding maps (Figure 3b, Figure **S2**). Quantitatively, neighborhood-informed decoding significantly reduces the posterior position standard deviation in all one perturbed maternal signal mutants (Figure 3d,e, Figure **S2**).

Together, these results suggest that neighborhood-informed position decoding may better characterize the underlying mechanism mapping gap gene expression to positional readout in the form of pair-rule gene expression.

## III. Discussion

In many contexts, cells acquire distinct position-dependent identities early in embryonic development [8, 9, 21, 22]. Therefore, many times, cells need to “know” their position in order to determine their future identity. Information theory provides a useful method to quantify the amount of information each cell has with regards to its position. Previous works using this approach have shown that while individual cells encode a substantial portion of the information necessary for unique identity specification, the embryo as a whole encodes the remainder [9]. However, it remained unclear where this information resides or how cells can access it.

To account for the gap in our understanding of unique cell identification, we looked into positional information embedded in the local environment of cells. This direction is inspired by a growing body of work showing that the local cellular neighborhood can strongly impacts cell fate [23–25]. We mathematically modeled these local influences and showed that they can provide significant positional information, enough to determine cell positions uniquely in realistic settings.

We further corroborated our theoretical results through empirical observations in *Drosophila* embryos. Specifically, in wild-type embryos, the minimal variance estimator using neighboring expressions is sufficient to close the positional information gap. Furthermore, we showed that integration of gene expression levels in the local neighborhood reduces ambiguity in the predicted position. This lower ambiguity then improves the prediction of pair-rule gene expression patterns downstream.

To test the extent to which positional readout is encoded in local gap gene expression, we analyzed mutant embryos with disrupted maternal signals. Although these embryos have altered gap gene expression and pair-rule stripe patterns, local gap gene expression levels can be used to predict cell position with significantly reduced ambiguity. We believe that the improvements are moderate in mutant backgrounds, because there may be additional factors that influence predicted pair-rule expressions. These findings suggest that positional information alone, given a wild-type optimal decoder, may not suffice as a model for a complete understanding of decoding under disrupted conditions, and can be further adjusted in future studies.

While our study focuses on a single axis in a relatively simple organism, it lays the groundwork for applying neighborhood-informed decoding to more complex systems. Future research could extend this methodology to multidimensional contexts and organisms with greater developmental complexity, advancing our understanding of how local interactions contribute to cellular identity and development.

## IV. Methods

### A. Decoding Position Given Neighboring Gene Expression

We will study the case where *g*_*n*_ represent the gene expressions at *x*’s adjacent positions, that is *g*_*n*_ = [*g*(*x* − 1), *g*(*x* + 1)]^*T*^. *g*_*n*_ ∈ **R**^2*d*^ and *g*_env_ = [*g, g*_*n*_] ∈ **R**^3*d*^ where *d* is the number of genes considered. Following previous work [8, 10], we will assume that the fluctuations in expression at each position along the AP axis are Gaussian, so that:

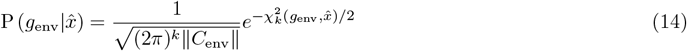

is the likelihood of the expression pattern *g*_env_ at position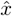, and:

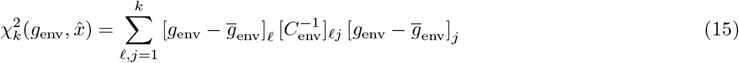

measures how similar the gene expression patterns in one embryo at position *x, g*_env_, is to the mean expression over embryos at the same position, 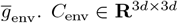 is the covariance matrix of *g*_env_.

Using Bayes’ law we can determine the posterior distribution over positions 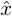 given the local gene expression, *g*_env_:

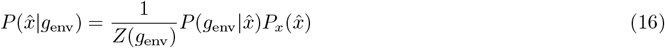

Here, 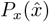 is the prior distribution that a cell is at position 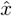, which, given no additional information, will be assumed to be uniform. *Z*(*g*_env_) serves to normalize the distribution and is independent of the position 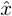.

### B. Decoding Map Representation

The decoding map is a probabilistic representation of the predicted positions relative to the ground truth positions, which is schematically described in Figure 1a. The x-axis represents the ground truth positions along the anterior-posterior axis, *x*. The y-axis represents the predicted positions 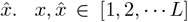 where *L* is the number of positions along the AP axis. Therefore, each column represents the posterior distributions over positions given gap gene expression patterns, namely 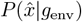 when decoding in a neighborhood-informed fashion, or 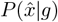, when decoding in a cell-independent manner. The decoding map is calculated per embryo, and we present the sum of the decoding maps over all embryos in logarithm scale, to retain a meaningful dynamic range.

### C. Quantifying the Error in Positional Prediction

Here we aim to quantify the accuracy of position prediction. To do so, we first compute the maximum a-posteriori (MAP) of the distributions 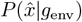 (neighborhood-informed) and 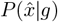 (cell-independent), as the predicted location given gene expression levels at the neighborhood environment of *x* or at the position *x*, respectively:

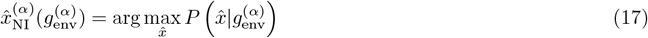

where 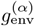 is the gene expression of cell *x* and its adjacent positions in embryo *α*. Together, 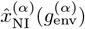 can considered as the neighborhood-informed (NI) position prediction. Analogously, for cell-independent decoding, the predicted position is given by:

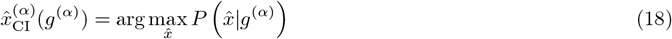

For each embryo *α*, we then calculate a vector of the MAP positions at each position along the AP axis:

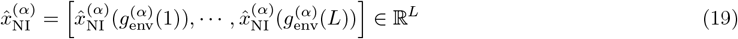

where *L* is the length of the embryo. The positions 1, …, *L* are the assumed ground truth positions, which we also arrange in a vector *x*_gt_ = [1, …, *L*] ∈ ℝ^*L*^. We define the analogous MAP vector for cell-independent decoding, 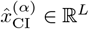

To quantitatively measure the positional prediction error, we compute the mean absolute difference between the ground-truth position and the MAP of the posterior distribution of predicted position over all embryos and over all positions. Specifically, the neighbor-informed positional prediction error is:

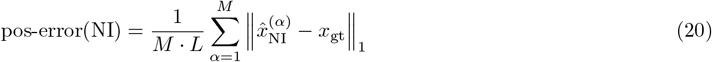

and the cell-independent positional prediction error is:

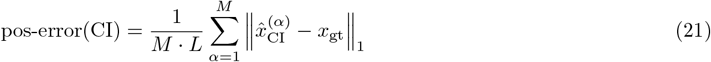

### D. Position Estimate Error

The position estimate error as defined in Equation 8 is explicitly calculated as [10]:

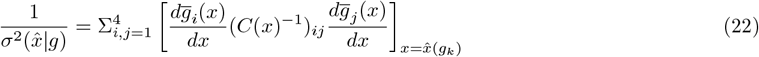

where *g*_*i*_ represents the expression of gap gene *i* (the gap genes include {Kr, Kni, Hb, Gt}), 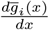 is the derivative of the mean expression over embryos of *g*_*i*_ at position *x*, and *C*(*x*) is the covariance matrix of the gene expression of the four gap genes at position *x*.

When decoding position with neighboring gene expression, the position estimate error 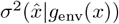 integrates the expression at the adjacent right and left positions, explicitly :

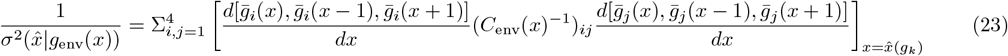

where *C*_env_(*x*) is the covariance matrix of the gene expression of the four gap genes at positions *x, x* − 1, and *x* + 1.

### E. Pair-Rule Gene Expression Comparisons and Reconstruction

Using the position decoder, we reconstructed the predicted wild-type pair-rule gene expression profiles along the AP axis of the embryo. This reconstruction of expression can be compared to the wild-type’s mean pair-rule gene expression profile across the AP axis, weighted by the noise in expression at each position. Formally, the expected expression of pair-rule gene *g*_*k*_ based on a neighborhood-informed decoding is:

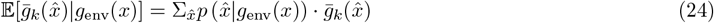

The analogous expression for a cell-independent position decoding:

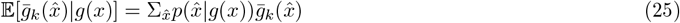

Where 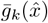 is the mean expression of pair-rule gene *k* in wild-type embryos at position 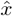.

The reconstruction error is then the absolute error between the ground-truth and predicted pair-rule gene expression. The neighborhood-informed reconstruction error is given by:

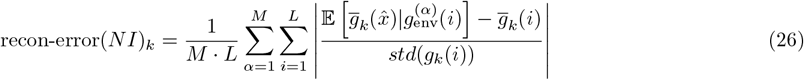

where *M* is the number of embryos, *L* is the number of positions along the AP axis, and *std*(*g*_*k*_(*i*)) is the standard deviation across embryos in gene expression of pair-rule *g*_*k*_ at position *i*. The reconstruction error for the cell-independent decoder is calculated analogously.

For mutant embryos, the ground truth expression of pair-rule genes, 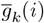, is taken for the mutant embryos and not the wild-type, however the reconstruction is done using the 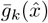 as in the wild-type case (Equations 24, 25).

## V. Data availability

The wild-type and mutant background *Drosophila* melanogaster datasets that we analyzed in this manuscript were collected in [8] and included in its Supplementary Materials.

## VI. Code availability

The code is publicly available via GitHub at https://github.com/nitzanlab/Neighborhood-Informed-Positional-Information.

## VII. Author Contributions

M.E, R.F and M.N. conceived the study; M.E. and R.F. developed the framework under the guidance of M.N. M.E performed the biological data analysis and implemented the software. R.F and M.E performed the analytical analysis. All authors interpreted the results, shaped the research, and wrote the manuscript.

## Supplementary Information

### 1. Data description

The empirical analyses were conducted based on data of *Drosophila* embryos collected in Petkova *et al*… For completeness, below we will shortly describe the experimental setup [8].

WT *Drosophila* embryos were stained fluorescently against the four trunk gap genes, Gt, Kr, Kni, and Hb simultaneously. Two sets of embryos were collected during nuclear cycle 14: *n* = 38 in the 40-44 minute time window, and *n* = 254 in the 38-48 minute time window. Analysis of WT pair-rule gene expression was conducted on *n* = 34 embryos, in the 45-55 time window, simultaneously stained for the *eve, prd*, and *run* pair-rule genes. The embryo fixation, antibody staining, imaging, and profile extraction were performed based on [26].

Mutant *Drosophila* embryos were created by perturbing the maternal signals Bicoid (Bcd), Nanos (Nos), and Torso-like (Tsl). Specifically, six mutant backgrounds were created by perturbing either one or two of these signals. We show results for the following mutant backgrounds: *etsl*^4^ (TOR-), *bcd*^*E*1^ (BCD-), *osk*^166^ (NOS-), *bcd*^*E*2^*osk*^166^ (TOR+), *nos*^*BN*^ *tsl*^1^ (BCD+), and *bcd*^*E*1^*etsl*^1^ (NOS+). The scaling of individual profiles was conducted as previously described in [3, 26]. The gap gene expression in mutant backgrounds was analyzed in the 38-48 minute time window for *n* = 40 *etsl*^4^ embryos, *n* = 20 *bcd*^*E*1^ embryos, *n* = 28 *osk*^166^ embryos, *n* = 15 *bcd*^*E*2^*osk*^166^ embryos, *n* = 19 Bcd-only germline close embryos, and *n* = 31 *bcd*^*E*1^*etsl*^1^ embryos. The pair-rule gene expression in mutant backgrounds was analyzed in the 45-55 minute time window for *n* = 14 *etsl*^4^ embryos, *n* = 12 *bcd*^*E*1^ embryos, *n* = 11 *osk*^166^ embryos, *n* = 17 *bcd*^*E*2^*nos*^*BN*^ embryos, *n* = 32 Bcd-only germline clone embryos, and *n* = 20 *bcd*^*E*1^*etsl*^1^ embryos.

### 2. Pairwise Correlations Close the Information Gap

In previous work, McGough *et al*. modeled the correlations in gene expressions between positions as:

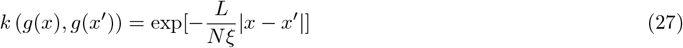

where *g*(*x*) is the expression of a specific gene at position *x, ξ* > 0 is the correlation length, *L* is the length of the AP axis in the embryo and *N* the number of positions along the axis for which the analysis is performed. If, in addition to the above, we also assume that the gene expression per position *g*(*x*) are Gaussian as in Methods Section A, then for every pair of positions the joint distribution *p* (*g*(*x* + 1), *g*(*x*)) is also Gaussian. The working assumption that the gene expression in pairs of positions are jointly Gaussian matches those made in previous work [9]. We will now proceed to show that one manner in which such a correlation structure can come about is if the gene expressions are Markovian along the AP axis. In other words, such correlation patterns arise when we assume that:

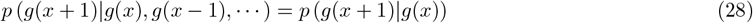

This Markovian structure will allow us to define the long-range correlations in terms of local correlations between adjacent positions.

For positions that are equally spaced along the AP axis, the above assumptions dictate the following Markov chain:

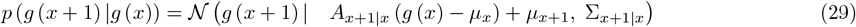

where *A*_*x*+1|*x*_ is the transition matrix of the Markov chain from *g*(*x*) to *g*(*x* + 1), and Σ_*x*+1|*x*_ is the covariance of *p* (*g*(*x* + 1)|*g*(*x*)). Since the joint distribution *p*(*g*(*x* + 1), *g*(*x*)) is a Gaussian distribution (through the assumptions above), then both *A*_*x*+1|*x*_ and Σ_*x*+1|*x*_ can be written in closed form as:

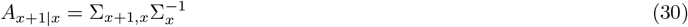

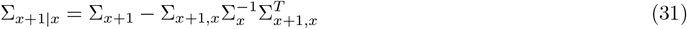

where 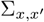 is the joint covariance of the gene expression *g*(*x*) and *g*(*x*^′^) at positions *x* and *x*^′^, respectively. Σ_*x*_ is the covariance of gene expression *g*(*x*) at position *x*.

Focusing on the correlations between gene expressions in different positions, we will next set the mean gene expression at position *x, µ*_*x*_ = 0, ∀*x* to simplify the following derivations. Additionally, we will assume for simplicity that the joint covariance matrix is constant throughout the chain so that Σ_*x*+1,*x*_ = *C* and Σ_*x*_ = Σ_*c*_ for any *x*. This, in turn, means that the transition matrix *A*_*x*+1|*x*_ = *A* is constant throughout the chain. Every step in the chain is then given by:

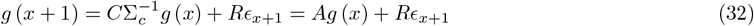

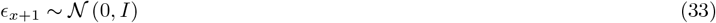

where we denoted the conditional covariance through its Cholesky decomposition, Σ_*x*+1|*x*_ = *RR*^*T*^ and where *ϵ*_*x*+1_ is the randomness introduced when transitioning from *g*(*x*) to *g*(*x* + 1).

We can now unroll the Markov chain using the step from Equation 32, up to Δ*x* steps:

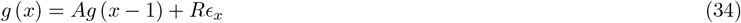

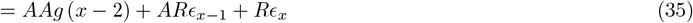

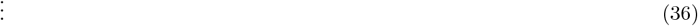

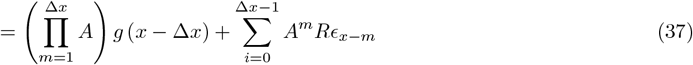

where we used the shorthand 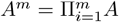. This allows us to explicitly write the covariance matrix of *g*(*x*) given the observation of *g*(*x* − Δ*x*):

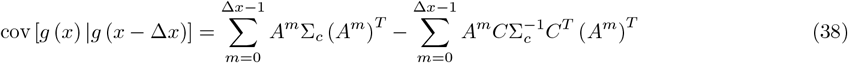

Using the properties of the Gaussian distribution, we can derive the joint covariance from the conditional covariance Σ_*x*|*x*−Δ*x*_ together with that of the marginal Σ_*x*−Δ*x*_. Additionally, we use the fact that the right hand side of Equation 38 has telescoping sums. All together, we get:

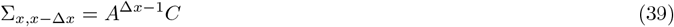

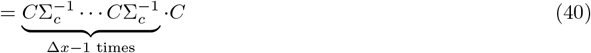

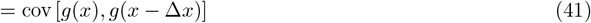

This expression is difficult to parse in the general setting, but can be better understood when we assume that the expression of one gene, *g*_*i*_(*x*), is independent of that of other genes, *g*_*j*_(*x*) ∀*j ≠ i*. This implies that 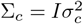 (where 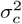 is the variance in expression of each gene) and the matrix *C* are both diagonal matrices. As such, we can rewrite *C* = diag (*c*_1_, …, *c*_*d*_) where *d* is the number of genes. In this case, we can analyze each gene independently, and we get:

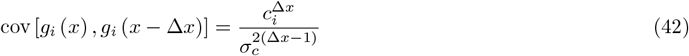

Furthermore, to relate to previous results [9], the correlations are given by:

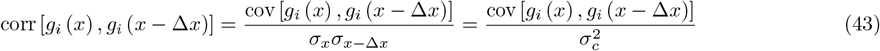

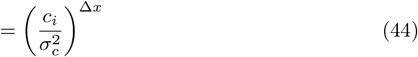

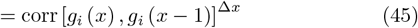

In the special case where 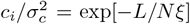 for some *ξ*, which is a reparametrization which is always possible whenever *c*_*i*_ > 0, we arrive at:

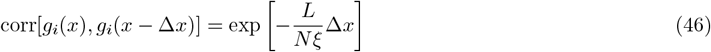

which is the same as Equation 27.

### 3. Decoding with Neighboring Gene Information Strictly Decreases Position Error

In this section we will show that the lower-bound on the amount of positional information strictly increases when adding neighboring information, as long as the correlations between the neighbors are not equal to ±1.

We will look at the case where the gene expression of *x, g*, and its neighbors *g*_*n*_ are jointly observed. Here we assume that *g* ∈ ℝ^*d*^ and *g*_*n*_ ∈ ℝ^*n·d*^ where *n* are the number of neighbors. A lower bound for the variance of any predictor 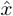 can be found using the Cramer-Rao bound. When only *g* is observed, this is equal to:

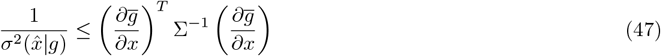

where 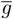 is the mean gene expression and Σ is the covariance matrix between all genes, which we will assume is independent of the position *x*. This is the cell-independent version of the Cramer-Rao bound, to which we will relate the neighborhood-informed version.

The Cramer-Rao bound for the neighborhood-informed decoder will depend on the inverse covariance between *g* and *g*_*n*_. As such, to explicitly write down the neighborhood-informed lower bound, we need to invert the covariance of *g* and *g*_*n*_:

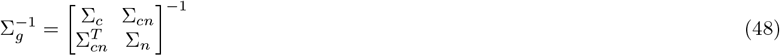

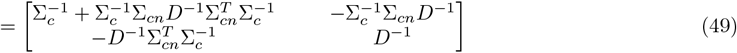

where Σ_*c*_ is the marginal covariance of *g*, Σ_*n*_ is the marginal covariance of *g*_*n*_ and 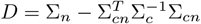.

Defining:

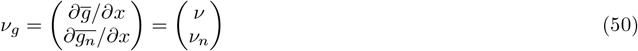

for ease of notation, the neighborhood-informed bound takes the following form:

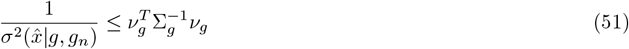

We can now plug Equation 49 into the above to get a precise term for the Cramer-Rao bound:

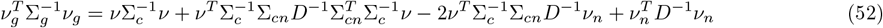

For ease of notation, we will define 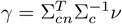. Using this shorthand, the above simplifies to:

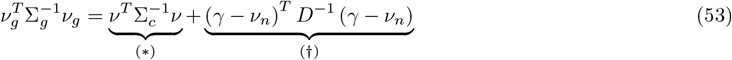

The (∗) is exactly the same as the cell-independent Cramer-Rao bound, the RHS of Equation 3.

To show that the neighborhood-informed bound on the variance is lower than that of the cell-independent, we must show that the (†) term is a positive number. As long as Σ_*c*_, Σ_*g*_ are both positive-definite, then *D* must also be positive-definite as it is Schur’s complement of Σ_*c*_ in Σ_*g*_. Because of this, the (†) term is also guaranteed to be positive. In that case:

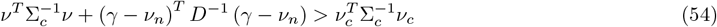

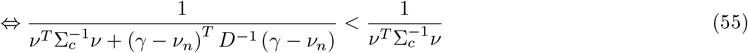

That is, the lowest possible variance 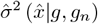 that can be attained when using information from neighbors is smaller than when not using neighbors:

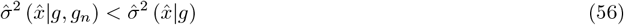

To summarize this relation, we will define:

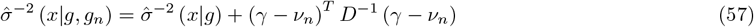

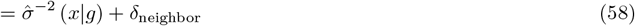

where *δ*_neighbor_ = (*γ* − *ν*_*n*_)^*T*^ *D*^−1^ (*γ* − *ν*_*n*_) is the added information from the neighbors.

### 4. The Cramer-Rao Bound is Reached Under Gaussian Assumptions

In the special case when it is assumed that *x, g, g*_*n*_ are jointly Gaussian, a closed-form expression for the variance of the decoder, 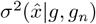, can be derived. We begin by following previous work and follow the assumption that the likelihood *p*(*g, g*_*n*_ | *x*) is a Gaussian distribution. Furthermore, we will assume that the prior over positions is also a Gaussian distribution:

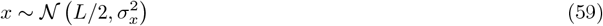

where *L* is the length of the embryo and 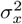 is the variance of this prior over the position, and will be taken to be much larger than the length of the AP axis.

The above Gaussian assumptions force a specific form on the conditional distribution *p*(*g, g*_*n*_ *x*), through the definition of conditional distributions of a Gaussian. In particular, the mean and covariance of this conditional distribution take the following form:

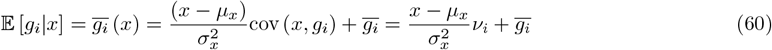

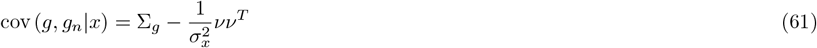

The value *ν*_*i*_ here is equal to cov (*x, g*_*i*_):

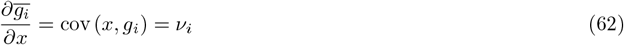

and is the same value as we had for the Cramer-Rao derivation in Equation 50, when the gene expression *g*_*i*_ is Gaussian conditional on the position *x*.

The log of the posterior distribution is given by:

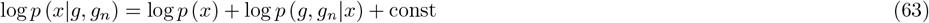

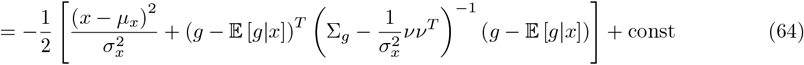

The Hessian of this log-posterior distribution will be the negative inverse of the variance, and is given by:

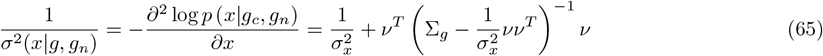

The matrix inverse in the above expression can be simplified using the matrix inversion lemma:

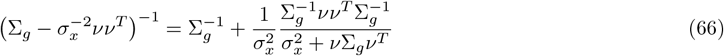

Using this simplified form of the inverse of the matrix, the variance is equal to:

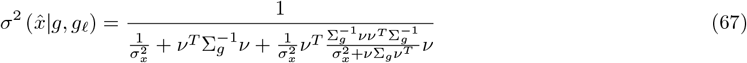

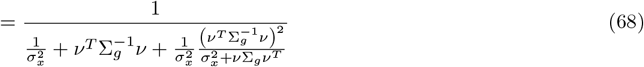

In our definition of the prior over the position *x*, it is centered around a specific point and approaches a uniform distribution when 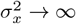. Taking this limit, we get the following:

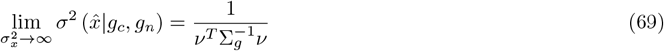

This is the same as the Cramer-Rao bound in Equation 3.

We can now use the fact that we have the explicit form of the Cramer-Rao bound in Equation 57. This allows to derive an exact expression for the information gap of the neighborhood-informed decoder under the Gaussian assumption:

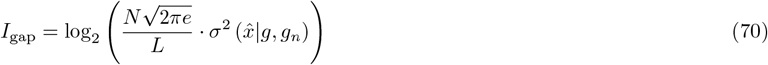

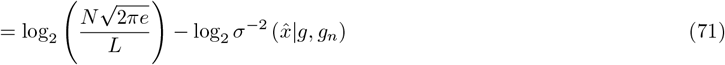

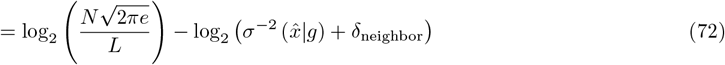

## Supplementary Figures

**Fig. S1:**
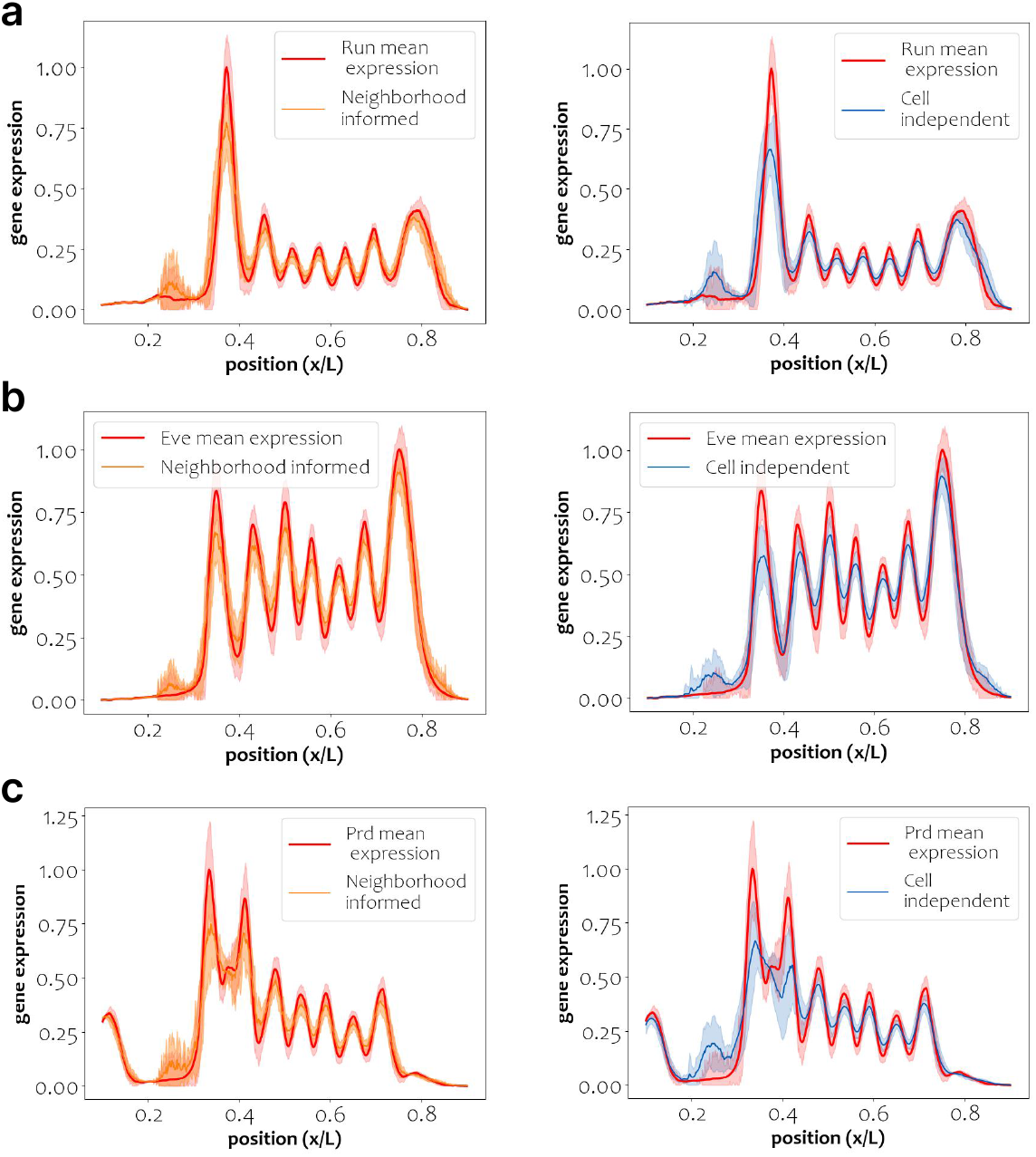
WT pair-rule gene expression predictions. The reconstruction of pair-rule gene expression spatial profiles based on the neighborhood-informed (orange) and cell-independent (blue) decoders, relative to the ground-truth mean pair-rule gene expression (red), for *(a)* Run, *(b)* Eve, and *(c)* Prd (also shown in Figure 2b).

**Fig. S2:**
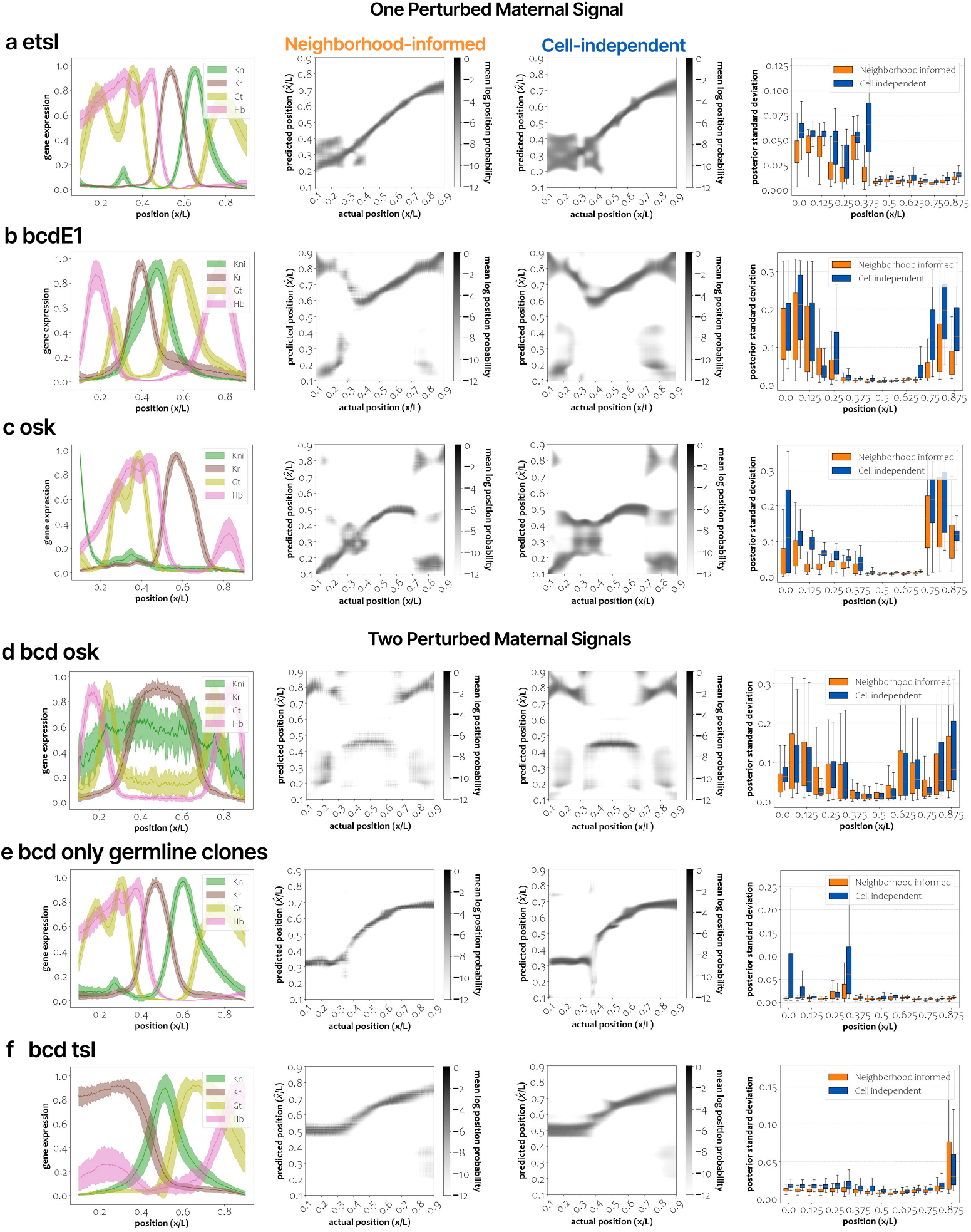
Position decoding and pair-rule stripe reconstruction in all mutant background embryos. Each row encases results for six types of mutant background embryos. The left most column shows gap gene expression profiles along the AP axis for each mutant type. The middle two columns show the decoding maps based on neighborhood-informed (left) and cell-independent (right) decoders. The right most column shows the posterior standard deviation over pair-rule expression reconstruction for the neighborhood-informed (orange) and cell-independent (blue) decoders, binned to 20 bins across the AP axis. Results are shown for the following mutants: *(a)* etsl, *(b)* bcdE1, *(c)* osk, *(d)* bcd osk, *(e)* bcd only germline clones, and *(f)* bcd tsl. The posterior position distribution standard deviation is significantly reduced when decoding in a neighborhood-informed manner (t-test p-value *<* 0.05) in all mutants with a single perturbed maternal signal (etsl, bcdE1, and osk) and in most instances of double perturbed maternal signal (100% of bcd only germline clones, 80% of bcd osk, and 40% of bcd tsl).

**Fig. S3:**
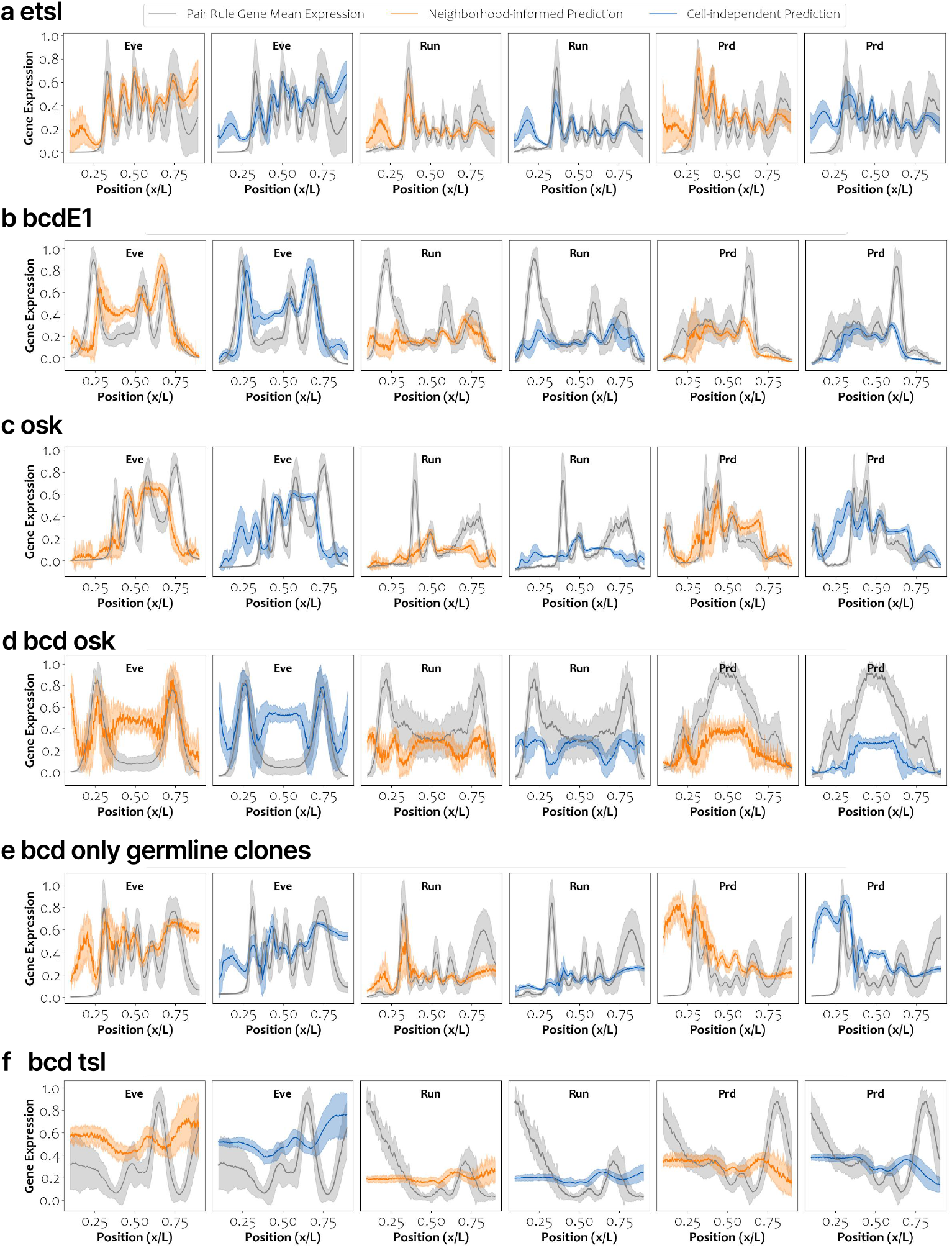
Pair-rule gene expression predictions in mutant background embryos. Each row presents the reconstruction of pair-rule gene expression spatial profiles based on the neighborhood-informed (orange) and cell-independent (blue) decoders, relative to the ground-truth mean pair-rule gene expression (gray), for Eve (first two columns), Run (next two columns), and Prd (last two columns), for each mutant type (rows).

**Fig. S4:**
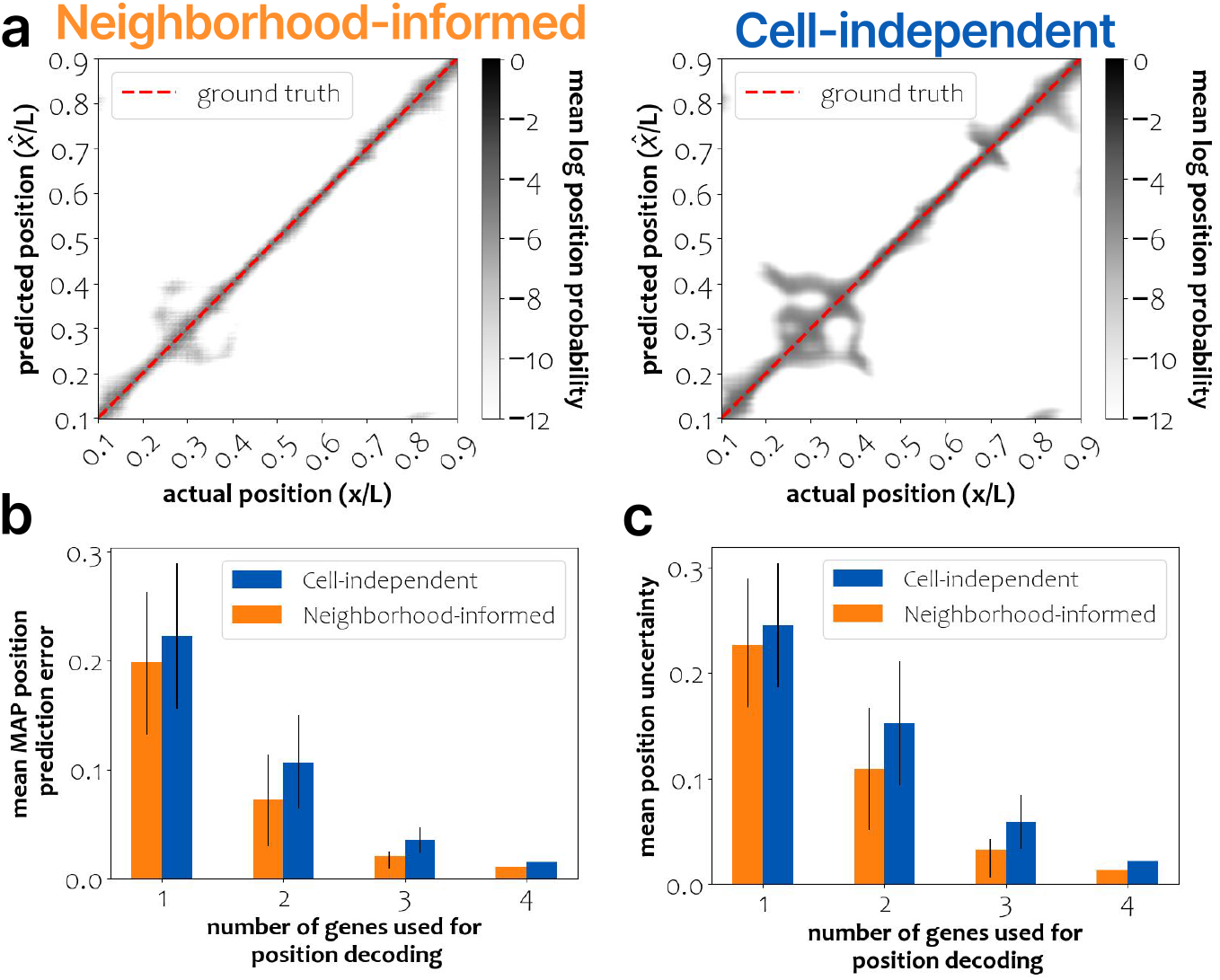
Position Decoding when training and testing on the same WT *Drosophila* embryo sample. The dataset of 38 WT embryos was used to infer the likelihood distribution and predict the positional posterior distribution. *(a)* Position decoding maps based on neighborhood-informed (left) and cell-independent (right) decoding given Kr, Gt, and Hb expression. (b)The mean error of position prediction using the MAP estimate, grouped by the number of gap genes used for decoding. For each group, we compare neighborhood-informed to cell-independent decoding. The predicted error is significantly lower for neighborhood-informed decoding (t-test p-value *<* 0.05 for embryo-to-embryo comparisons) for all gene subsets. *(c)* The standard deviation of the predicted position distribution, grouped by the number of gap genes used for decoding, is significantly lower for neighborhood-informed decoding (t-test p-value *<* 0.05 for embryo-to-embryo comparisons) for all gene subsets.

